# Highly Sensitive and Specific Amyloid-β Oligomers Detection Using CRISPR-Cas12a Probe System aided by Graphene Oxide for Early Diagnosis of Alzheimer’s Diseases

**DOI:** 10.1101/2022.09.27.509482

**Authors:** Jiaqi Tang, Zongwei Zhu, Chuanxu Yang

## Abstract

Alzheimer’s disease (AD) is highly age-specific that results in cognitive incapability including dementia. It is proposed that AD is caused by the dysfunctional gene producing excessive amyloid-β protein that eventually results in aggregated form of amyloid-β Oligomers (AβO). We couple the CRISPR-Cas12a system with a specially designed single-stranded DNA (ssDNA) aptamer that specifically forms a hybrid complex with AβO. We exploit the aptamer as the substrate for the crRNA-aptamer duplex due to its high flexibility, sensitivity, and selectivity. We further adopted the graphene oxide (GO) fluorescent system to couple with our CRISPR-Cas12a probe system. Our coupled probe system can effectively detect the presence of AβO by quantitatively returning fluorescent signals with high specificity and accuracy.

## 1. Introduction

Alzheimer’s disease (AD) is highly age-specific that results in cognitive incapability including dementia [1]. It is observed that the number of AD patients has been increasing in recent years across the world and is estimated to be 106.8 million in 2050 [2]. It was originally believed that the cause of AD was primarily due to the production of insoluble amyloid-β protein (A-β protein), namely A-β fibrils (AβF), that induces toxicity to neuron cells [3]–[5]. It is not until recently that a new hypothesis was proposed, which brought the amyloid-β protein oligomer (AβO) into the spotlight. It was found that soluble AβO was more strongly correlated with AD onset than AβF does. The accumulation of abnormally folded A-β protein, caused by the discrepancy between the rate of production and clearance, is believed to be the cause of the development of neurofibril tangles and the primary driver for AD-related pathogenesis [6]–[8]. More detailed pathogenesis was investigated in the new pathological studies; AβO was suspected to induce both biological and physiological toxicities to neuron cells, including the blockage of ion flow, synaptic transmission, and damage to kinase activities, that eventually leads to the long-term, irreversible impairment of memory retrieval and other cognitive abilities [3], [9]–[11].

Early diagnostics and therapeutics are important factors to treat Alzheimer’s disease [1]. For example, the vascular hypothesis is a key step to the preclinical prediction of dementia using neuroimaging [12] and has been widely studied in recent years [13]. Current clinical assessment only provides approximately 85% accuracy based on neuroimaging, personal history, and cognitive assessment [3], [14], [15]. In addition, researchers have developed nonpharmacological treatment for AD via the mind-brain approach [15]–[17]. On the other hand, it has been shown that visual memory could predict AD much earlier than diagnosis [14]. It was found that the E22G pathogenic mutation of β-amyloid can enhance the misfolding of Aβ1-40[18]. Other approaches to treating Alzheimer’s disease include the amyloid cascade hypothesis [4] as well as the monoclonal antibody with conformational specificity for a toxic conformer [5].

It is important to note that the initial symptom of AD could occur many years after the onset of the disease [5, 15], so the aforementioned diagnostic methods not only suffer from high cost, high complexity, and low accuracy, but also their application is severely limited due to high operation cost, complex technical procedures, personal biases, and late-informative stages. Early diagnosis is often difficult but can provide patients with a more distinct explanation for the cause and greatly facilitate effective treatment at the same time reducing financial cost [19]. Therefore, an urgent need to develop an early, accurate, and cost-efficient diagnostic method is present.

Clustered, regularly interspaced, short palindromic repeats (CRISPR) Cas system is part of the immune system that many bacteria and archaea possess to withstand invasion and predation from plasmids and viruses [20]. CRISPR-Cas family has been exploited for its potential to act as a detection and diagnostic tool set due to its endonuclease activity. In this work, we particularly exploited the CRISPR-Cas12a due to its independence of initiating cleavage activity regardless of the maturation of crRNA and due to its featured indiscriminate ssDNase activity [21]. In this work, the initiator, crRNA, is a pre-determined ssDNA sequence, namely aptamer, that could 3D self-fold to form a hybrid with both biomolecules and the crRNA site in CRISPR-Cas12a. In addition, the *trans*-cleavage activity by CRISPR-Cas12a allows it to perform non-target DNA degradation. In this work, we coupled the FAM and GO as the fluorescence reporter system. By coupling it with the fluorescence system, CRISPR-Cas12a allows for detectable fluorescent signal output. Nevertheless, the predecessor has limited their target at nucleotide bases while the detection at the molecular level is urgently and massively needed. We have improved upon the established model of detection and created an effective, innovative, and cost-efficient detection method targeting AD by tracing the presence of AβO.

Graphene oxide is a 2D carbon nanomaterial that possesses a unique structure containing abundant pi electrons, which enables it to display featured photoelectric and physisorption property that is ideal to serve as a biosensor tool [22], [23]. In addition, the carboxyl and hydroxyl groups allow GO to possess high aqueous solubility which expands its potential as a biosensor tool [24]. Due to fluorescence resonance energy transfer (FRET), GO can effectively quench the fluorescence emitted by the FAM unit with its abundant pi electron. GO has been shown to have physiological absorption towards ssDNA through hydrogen bonding and pi pi stacking [25], [26], and such bonding has been shown to display a higher affinity to Adenine [23]. Therefore, we specially designed the FAM by adding a poly-Adenine tail of 20 units (A20) onto the FAM to enhance the physisorption between the GO and the FAM-A20. Due to its effective and selective quenching ability, GO is an ideal tool to act as a switch that returns fluorescent signal output only upon the encounter of the target molecule.

We exploited the aptamer as the substrate for the crRNA-aptamer duplex due to its high flexibility, sensitivity, and selectivity [27]. The nature of the aptamer allows it to form complementary base pairing with sequence series, while such pairing can be easily disrupted by competitive interaction between the aptamer and the target [28]. Aptamer’s 3D self-folding structure allows it to selectively form a complex with a broad range of targets including other nucleic acids, metal ions, and proteins through hydrogen bonding, hydrophobic stacking, Van der Waals forces, etc [29]. Aptamer has been shown to possess especially high affinity and specificity with proteins [28]. In this work, aptamer acts as a crRNA-AβO interface that can form a hybrid with both crRNA and AβO. To ensure the selectivity and accuracy of our probe system, we specially designed the sequence order in which the aptamer was mixed with crRNA and AβO. The aptamer was mixed with AβO before it does with crRNA.

Here, we coupled CRISPR-Cas12a system with a specially designed ssDNA aptamer that forms a hybrid complex with AβO. To establish the accuracy and quantitative signal output, we exploited GO as a tool for manipulating the fluorescent signal intensity as part of the “quencher” in the FQ reporter system, which changes the level of fluorescence based on the concentration of AβO in the substrate. Due to the high affinity of GO to Pyridine, we added a Poly-A tail of 20 units onto the FAM fluorescent unit, establishing a hybrid of FAM-A20 to enhance the physisorption and ensure effective FRET. We adopted an artificially selected ssDNA aptamer that could selectively form a hybrid with AβO and crRNA. We demonstrated that our detection system could quantitatively detect the concentration of AβO using aptamer-activated CRISPR-Cas12a and FQ reporter system consisted of FAM-A20 and GO with great accuracy, efficiency, and selectivity.

## 2. Experimental section

### Preparation of AβO

The ssDNA aptamers have distinct binding affinities to AβO evaluated by enzyme-linked oligonucleotide assay. Next, Chemicals and Milli-Q water in the experiments rely on the analytical grade. Professor Tsukakoshi et al have prepared and described the AβO [5, 33]. First, lyophilized Aβ1-40 was treated with hexafluroisopropanol (HFIP) on ice. Next, the Milli-Q water was added to the lyophilized Aβ1-40 and remained around twenty minutes at the room temperature. Then the solution was centrifuged at 14,000 g for about 15 minutes. Next, the supernatant was related to Ar-gas bubbling to remove HFIP. Then, the solution was incubated for a day at 800 rpm by stirring and centrifuged at 14,000 g for 20 minutes. Next, the supernatant was concentrated by using a molecular weight cut-off filter to obtain the concentrated AβO solution. At last, the prepared AβO was measured on the concentration by UV/Vis spectrophotometer at 280 nm.

### Synthesis of CRISPR-Cas12a

First, 10 nM CRISPR-Cas12a (80 μL) was mixed with 10 nM crRNA (80 μL). Then, the mixture was incubated at 25°C for 30 min. Lastly, the solution was mixed with 400 nM FAM-A20 (80 μL) and incubated at 25°C for 5 min.

### Preparation for AβO Detection with Florescence

CRISPR-Cas12a-crRNA (80 μL) solution was mixed with AβO (80 μL) of different concentrations at 25°C for 60 min. Next, the reaction was stopped by incubating the mixture at 60°C for 5 min. Then, the solution was mixed with 20 μg/ml graphene oxide (80 μL) at 25°C for 30 min. Finally, the fluorescent activity was measured by spectrophotometer with the excitation point of 485 nm with the slit of 3.5 nm and emission point of 495-650 nm with the slit of 1.5 nm.

### Preparation of Buffer A

Buffer A was prepared by adding 40 mM 4-(2-hydroxyethyl)-1-piperazineethanesulfonic acid (HEPES), followed by 100 mM NaCl, and lastly by 20 mM MgCl_2_. The pH of Buffer A is 7.4.

### Preparation of GO

First, the GO powder (25 mg) was mixed with ultrapure water (25 ml). Next, the mixture was then ultrasonic dispersed at 300W for 30 min and centrifuged at 4000 RPM for 5 min. Lastly, the supernatant was stored at 25°C.

### Characterization of GO

1 μg/ml GO power was added into the ultrapure water and sufficiently dispersed. Next, the solution was ultrasonic dispersed for 25 min. Then, the solution was added dropwise to the mica isinglass and remained until the fully naturally vaporized. Finally, the mica isinglass was placed into the AFM for scanning, in tapping mode, at scan rate of 1 Hz, scan size 3μm, number of pixels 512 × 512, under ambient conditions [3].

### AβO aptamer sequence

5’-GCCTGTGGTGTTGGGGCGGGTGCG-3’

### FAM-A20 sequence

5’-FAM-AAAAAAAAAAAAAAAAAAAA-3’

### AβO sequence

DAEFRHDSGYEVHHQKLVFFAEDVG SNKGAIIGLMVGGVV

## 3. Results and discussion

**Scheme 1.**
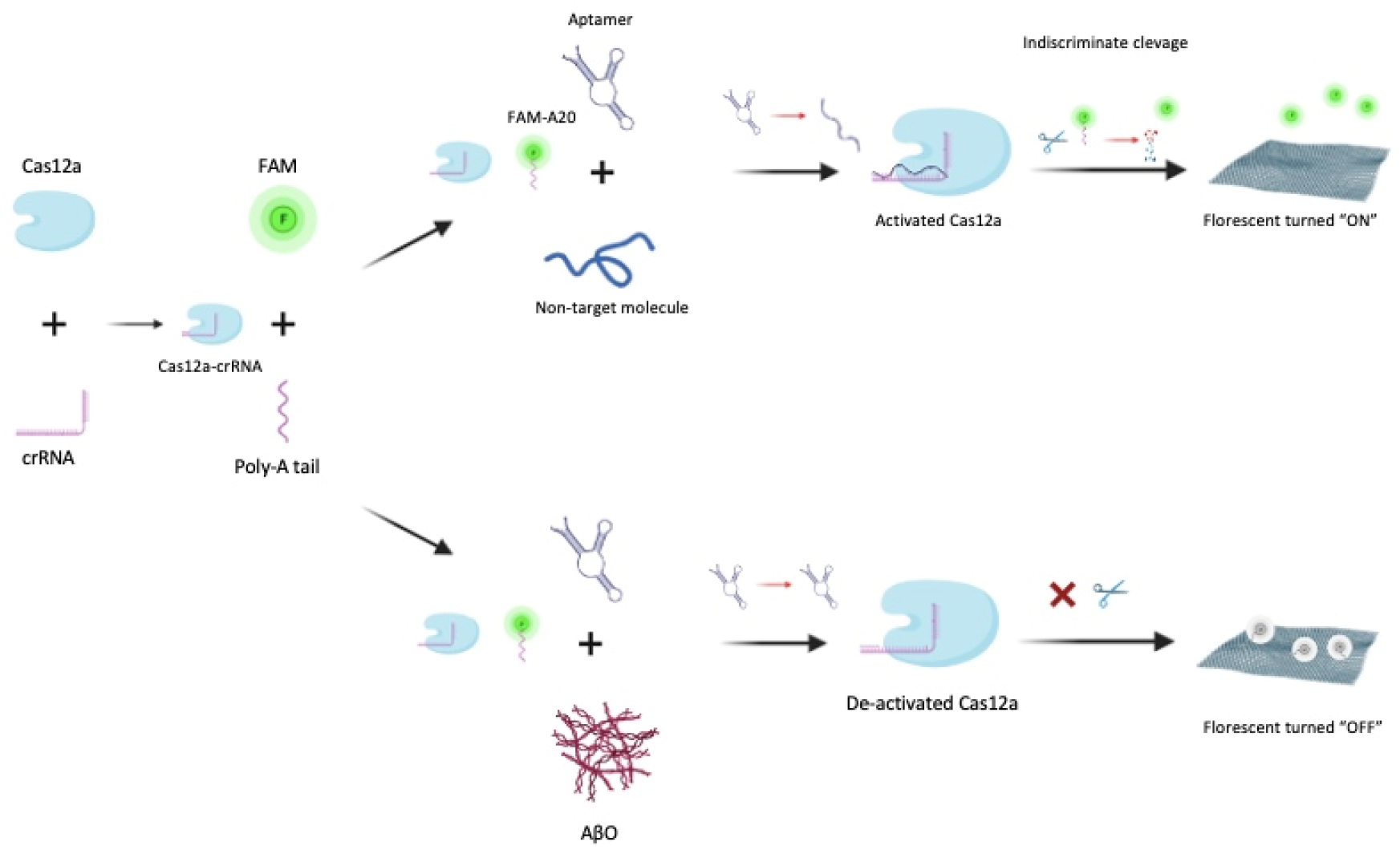
The working principle of the probe system

### Experimental working principle

First, CRISPR-Cas12a was mixed with crRNA to obtain the Cas12a-crRNA hybrid. Next, the Cas12a-crRNA hybrid was mixed with FAM-A20 reporter solution. Then, the Cas12a-crRNA-FAM-A20 hybrid was mixed with aptamer and target solution. Next, the hybrid was added to the GO solution. Last, the fluorescent spectrophotometric measurement was carried out. The 3D structure of aptamer undergoes conformational change when forming hybrid with AβO, which prevents it from further binding with crRNA in CRISPR-Cas12a protein. If the environment lacks AβO, then the free aptamer will activate CRISPR-Cas12a to initiate non-specific cleavage activity. Then, the Poly-A tail on the FAM-A20 will be cleavaged into small nucleotide fragments, which prevents it from absorbing onto GO to be quenched. Thus, one could receive positive fluorescent signal output. On the other hand, if the environment contains AβO, then aptamers are “occupied” by AβO and can not activate CRISPR-Cas12a. Thus, the FAM-A20 will not be cleavaged, which can be effectively absorbed and quenched by GO to produce the negative fluorescent signal output. Our coupled system can effectively and quantitatively detect the presence of AβO with high specificity and high accuracy using the fluorescence signal output (Scheme 1).

GO should be in single flake (single-layered) form in order to quench the FAM effectively [31]. Our available GO powder indicated that the diameter of GO is 1-1.2 μm as documented by the manufacturer. GO power (1 μg/ml) was dissolved into ultrapure water and performed ultrasonic dispersion to obtain the single-layered of GO to be used in our research. To confirm the form of the single-layered GO, AFM image of the sample was performed. AFM image (a) confirmed that the diameter of the GO is around 400 nM, suggesting the high effectiveness of removing GO with our method. The height profile (b) confirmed that the 3D height of our sample is at 1-1.2 nm, which is in accord with the GO morphology in the work by Jiang et al. [23]Next, we performed the dynamic light scattering to obtain a range and distribution of the size of the GOs in our sample. The graph suggests that the range of our sample shows great consistency and that most of the GO is around 500 nm, with a normal distribution curve. To improve the accuracy of our experiment, we investigated the interference of the concentration of GO to the fluorescent signal intensity in Buffer A, 20 μg/mL GO, and 4 μg/mL GO. We found that Buffer A has a very insignificant absorbance rate and that at 20 and 4 μg/ml GO, the absorbance range is at 200-850 nm. This graph confirms that the working concentration of our GO, which is 4 μg/mL, has very little absorbance rate that interferes with the fluorescent signal activity at 485 nm, suggesting the high feasibility of using 485 nm excitation wavelength as the working wavelength while guaranteeing the high accuracy.

Research has shown that the A-β protein will only induce AD when it is aggregated to form oligomer cluster from the peptide chain [32]. Therefore, to ensure the effectiveness of our research method, we obtained the A-β oligomer (AβO). AFM imaging was performed to confirm that A-β protein indeed aggregated and clustered to form into the 3D-shaped AβO with height in Figure 2. The heigh profile (b) suggests that the AβO in the sample has consistent height and 3D conformation, which is in accord with the established AβO 3D conformation [3], suggesting high effectiveness in our construction method. AFM imaging and height analysis was performed on the AβO 30 days after the construction (a) and (b), respectively. Both analyses show consistency with the AβO 30 days before, suggesting the insignificance of long-time storage time on the 3D conformation of the AβO.

**Figure 1.**
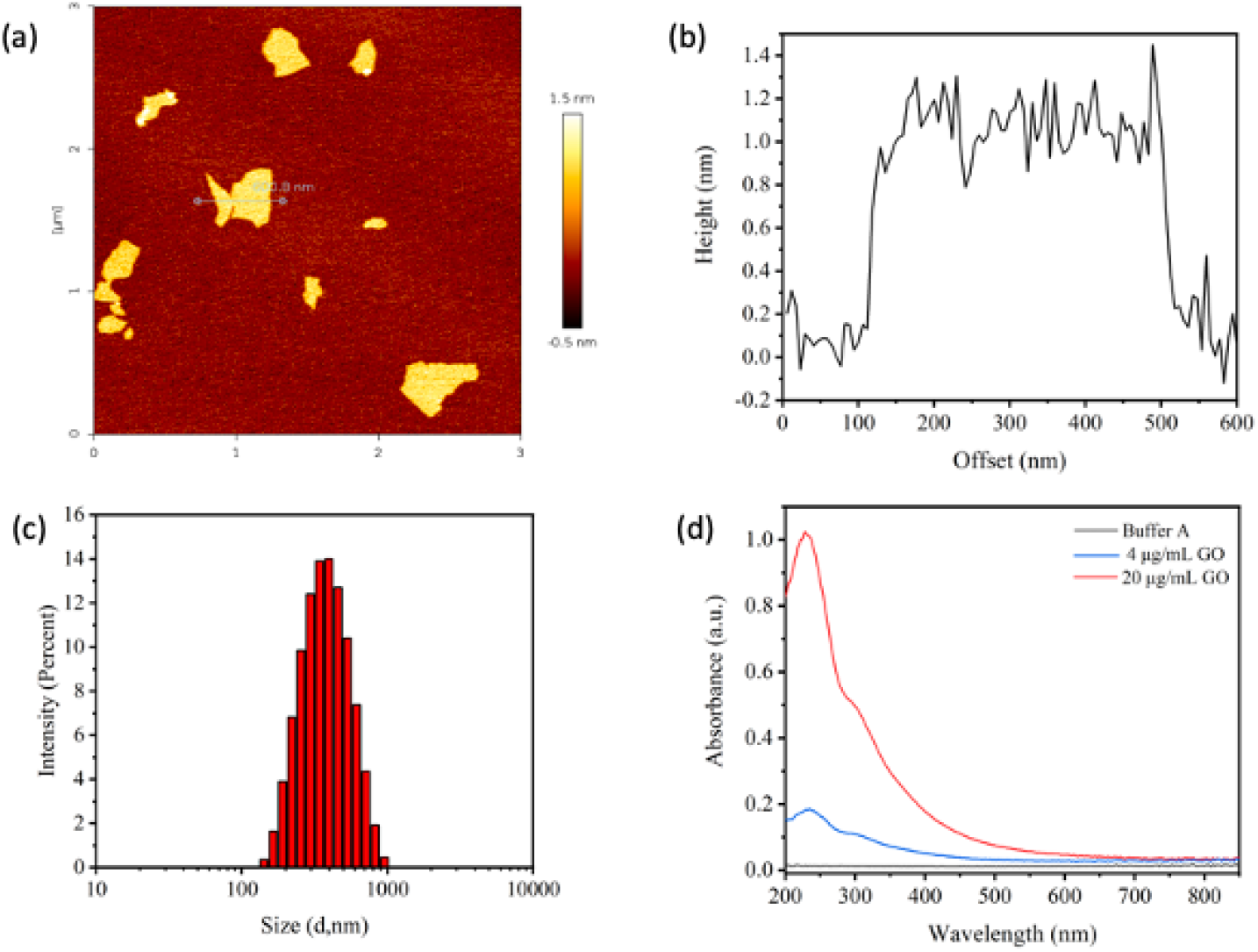
Characterization of GO. AFM image of GO (a). The height profile of GO (b). The size profile of GO was measured using dynamic light scattering (c). The absorbance activity of Buffer A, 4 μg/ml GO, and 20 μg/ml GO (d).

**Figure 2.**
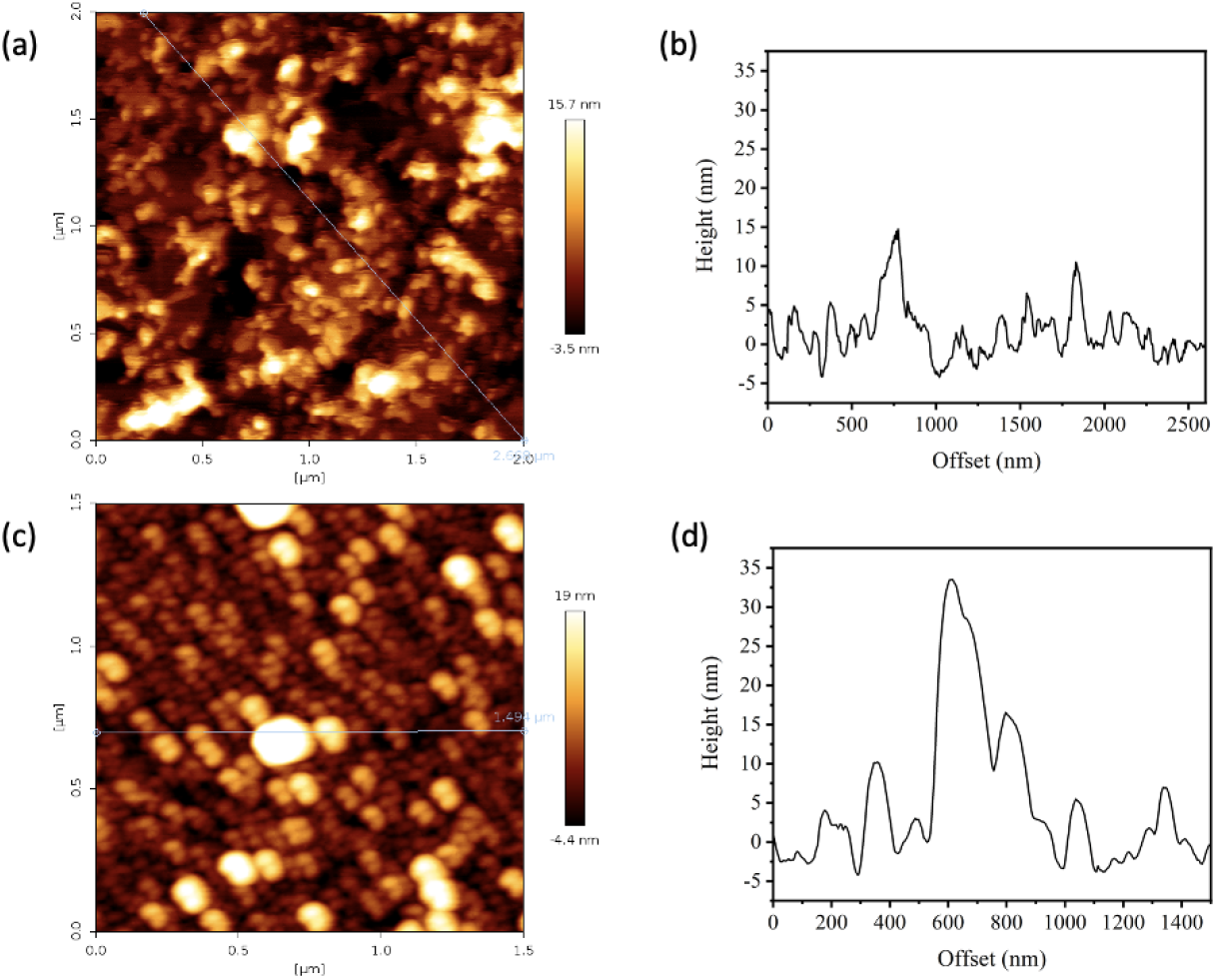
Construction of AβO. AFM image (a) and height profile of AβO on the day of construction (b). AFM image (c) and height profile of AβO 30 days after the construction (d).

We investigated the excitation and emission efficiency of the fluorescent spectrophotometer (Figure 3.) To guarantee the maximum and complete excitation spectra of the fluorescent signal, the excitation length is 485.0 nm. To ensure the optimal and maximum fluorescence signal output, the emission spectra are 520.0 nm. The fluorescent intensity was normalized for easier graph presentation, showing that the excitation and emission spectra are symmetrical and make the peak of both excitation and emission spectra more obvious and understandable.

**Figure 3.**
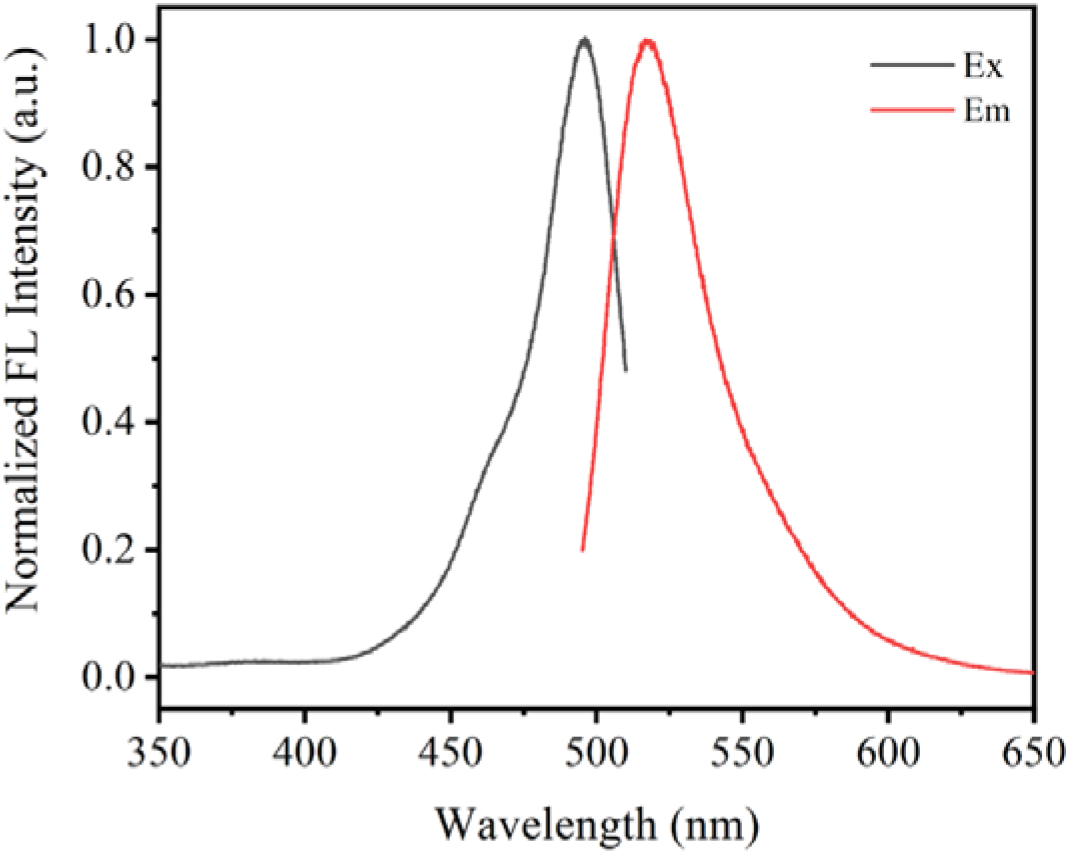
The excitation (Ex) and emission spectra (Em) of FAM

We investigated the quenching efficiency of 5 μg/mL GO to 100 nM FAM using GO concentration assays (0.0, 0.1, 0.5, 1.0, 2.5, 5.0, 10.0, 25.0, 50.0, 100.0 nM). As shown in Figure 4 (a), no significant fluorescence signals were observed at 100 nM GO at 520.0 nm, while the fluorescence signal peaked at 0 nM GO at 520.0 nm, which increased approximately 5000-fold (Figure 4 b). This suggests a well-established negative quantitative response and high sensitivity in our probe system. Figure 4 (b) suggests that the quenching efficiency grows logarithmically with the increase of GO concentration. The quenching efficiency is above 90% when the GO concentration is 5 nM. Since a higher concentration of GO will result in higher absorbance activity by GO (Figure 1 d) and that 90% quenching effect is sufficient to produce a reliable and accurate quantitative negative response, 5 μg/mL was adopted as the final working concentration.

**Figure 4.**
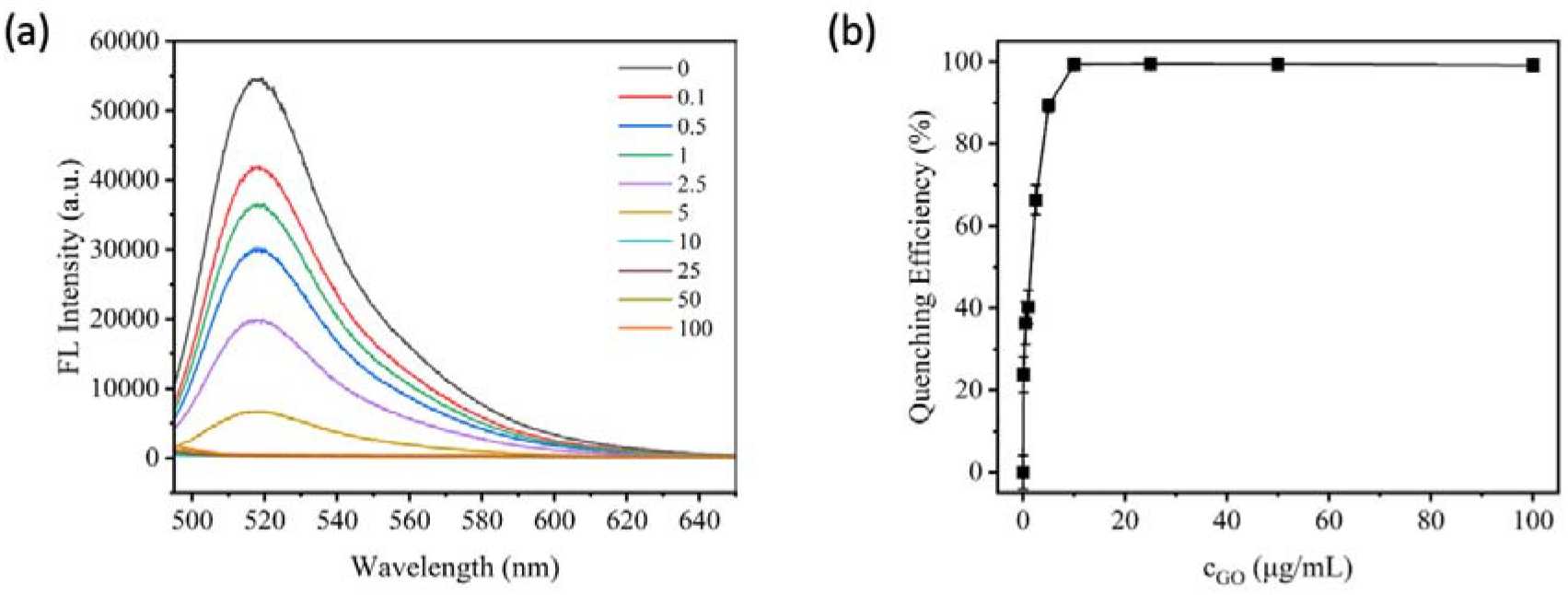
The quenching efficiency to 100 nM FAM of different concentration of 5 μg/mL GO (0.0, 0.1, 0.5, 1.0, 2.5, 5.0, 10.0, 25.0, 50.0, 100.0 nM)

We investigated the readability/sensitivity of the probe system under different FAM concentrations. The fluorescent intensity was modified into an logarithmic growth for better data presentation. The strongest intensity of fluorescent signal output was observed at 100 nM FAM, but the ratio between the fluorescent intensity with and without aptamer is too small which undermines its sensitivity. The ratio of the fluorescent intensity with and without aptamer peaks at 60 nM of FAM. However, the intensity of the fluorescence signal is not strong enough to ensure the best and most accurate measurement. Therefore, the concentration of FAM is decided to be 80 nM, in which the sensitivity is high while the fluorescence signal intensity is strong enough to be measured accurately and effectively (>5000 a.u.).

With the established optimal FAM-A20 concentration, we investigate the sensitivity and responsiveness of different concentrations of AβO to our probe. As Figure 5 (b) suggests, the fluorescent activity decreases with the increase in AβO concentration. The fluorescent intensity peaks at 0.05 nM of AβO, while reaching its lowest point at 50 nM. For a clearer graph presentation, we plotted the fluorescent intensity against the concentration of AβO using a logarithmic scale (Figure 6 a), which returns a strong negative logarithmic relationship between the two (R_2_ = 0.99019). This relationship suggests that our probe system can establish a strong and reliable quantitative measurement of AβO from the intensity of the fluorescence. thus proving a promising tool for medical and detection purposes. high accuracy and sensitivity and precision with established quantitative reliable ratio.

**Figure 5.**
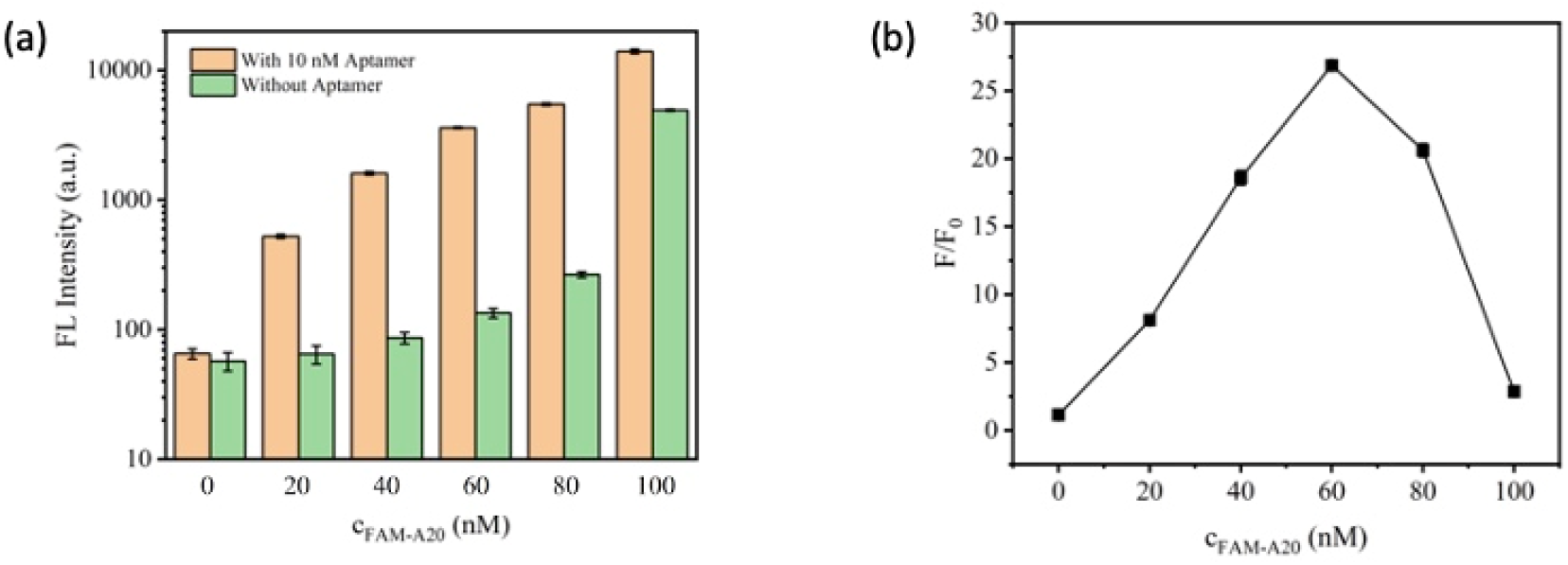
Fluorescent intensity of the FAM solution with and without aptamer (a). the ratio of the fluorescent intensity of the FAM solution with and without aptamer (b)

**Figure 6.**
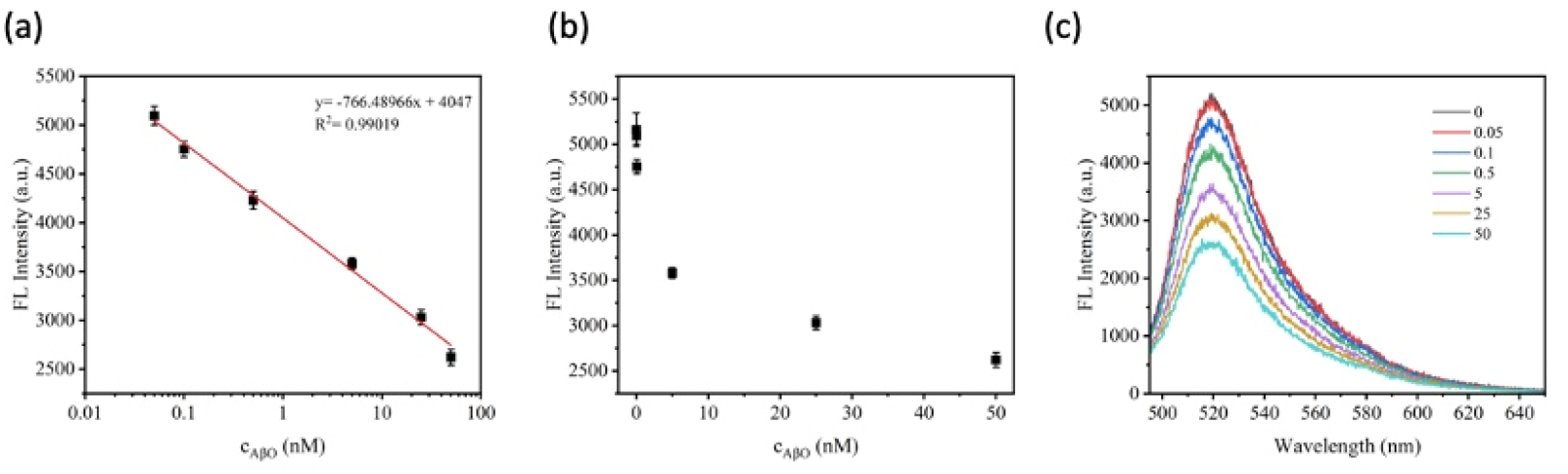
The fluorescent intensity of the probe based on different concentrations of AβO. The correlation analysis of between the concentration of AβO and the intensity of fluorescent intensity (a). The logarithmic relationship of the concentration of AβO to the fluorescent intensity (b). The wavelength and fluorescent intensity of different concentrations of AβO (c).

Lastly, to ensure the selectivity of our probe system, we investigated the response sensitivity of our probe to Aβ1-40 and AβO. Figure 7 (a) suggests a significant difference, more than double, in the fluorescent signal intensity between the Aβ1-40 and AβO. Figure 7

**Figure 7.**
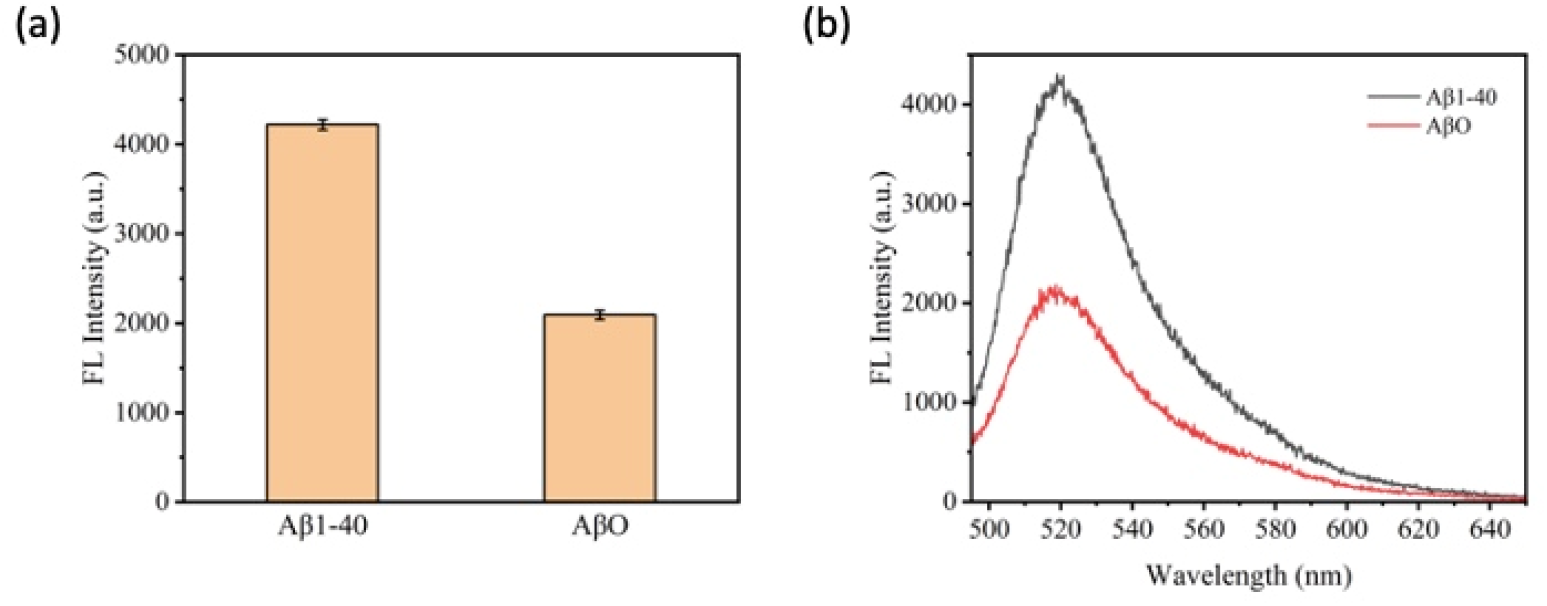
The fluorescent signal intensity of the probe with Aβ1-40 and AβO (a) and at different wavelengths (b).

(b) suggests that at our optimal emission spectra, 520.0 nm, the difference in the fluorescent intensity with Aβ1-40 and AβO shows the greatest difference. Both results suggests that our probe possess high selectivity that enables an effective and accurate diagnostic ability and power to AβO presence, thus indicating a promising future of our probe system for diagnostics.

## 4. Conclusion

This study has introduced the complex biosensing system consisting of AβO-specific aptamer as the target, FAM-A20 and GO as the reporter system, and CRISPR-Cas12a as the effector system. We have coupled the CRISPR-Cas12a system with a specially designed ssDNA aptamer that specifically form a hybrid complex with AβO. We also exploited the aptamer as the target for the crRNA due to its high flexibility, sensitivity, and selectivity. We further proposed to couple the adoption of GO system with our CRISPR-Cas12a detection system. We have successfully constructed an ideal 2D GO surface and 3D AβO conformation that is essential to real-life application, verified by AFM imaging. The most efficient excitation and emission wavelength were found to be 485.0 nm and 520.0 nm respectively. The quenching efficiency of GO peaked at 100 nM and reached the lowest point at 0 nM, returning an exponential relationship between the concentration of GO and its quenching ability. To ensure the readability of the probe system, the concentration of FAM-A20 was investigated to be at 80 nM. The sensitivity and quantitative substrate detection ability were investigated using different concentration of the target substrate AβO. The most intense fluorescent signal output was observed at 50 nM AβO, while the least intense fluorescent signal output was observed at 0.05 nM AβO. The AβO concentration and fluorescent intensity shows a strong negative logarithmic relationship, suggesting an ideal quantitative signal fluorescent signal output based on substrate concentration. Since only the aggregated form of A-β protein, namely AβO, has been shown to induce AD, we tested the responsiveness of our probe system to a non-target but highly similar molecule Aβ1-40 protein. The significant difference in fluorescent signal output suggests that our probe system has shown great selectivity and specificity towards the target AβO over non-target Aβ1-40. This work has shown that our coupled biosensing system can effectively detect the presence of AβO protein with high specificity and high accuracy.

**Table 1.** Comparison of CRISPR-Cas12a detection between this work and other probe system involving GO

| Probe | Methods | Linear range | Detection limit | Reference |
| --- | --- | --- | --- | --- |
| FAM-A20-GO | Fluorescence | 0.05-50 nM | ----- | This work |
| FAM-BHQ2-GO | Fluorescence | 0.5-150 nM | 0.44 nM | [23] |
| Cy5-ssDNA-GO | Fluorescence | ----- | 1.27 nM | [23], [33] |
| Cy5-MB-GO | Fluorescence | 0.1-10.0 nM | 0.03 nM | [23], [34] |
| TAMRA-DNA-GO | Fluorescence and Anisotropy Assay | 0.0-16.0 nM | 0.047 nM | [23], [35] |

